# Longitudinal characterization of the captive adult and tadpole Wyoming toad (*Anaxyrus baxteri*) microbiome

**DOI:** 10.1101/2023.06.13.544657

**Authors:** Nicole Scarberry, Zachary L. McAdams, Derek Benson, Jason Herrick, Brandon Moore, Aaron C. Ericsson

## Abstract

At one time thought to be extinct in the wild, the Wyoming toad (*Anaxyrus baxteri*) is one of the most critically endangered North American amphibian species. Despite approximately 20 years of *ex situ* breeding and reintroduction programs, these animals remain functionally extinct in the wild. There is concern among those working in these programs that individuals bred in captivity fail to develop the proper microbiome to withstand the stressors of their native habitat following release. In related species, the skin microbiome has been shown to have a defensive function against common pathogens affecting these animals. However, the early-life microbiome of developing tadpoles in this species remains unknown and therefore this defensive function is unexplored in the Wyoming toad. This study employed 16S rRNA amplicon sequencing to document the baseline microbiome of tadpoles bred for release and captive adult breeder populations. To characterize microbiome development, multiple rounds of skin mucosal and cloacal swabs were obtained concurrently from adult Wyoming toads bred at Omaha’s Henry Doorly Zoo and Aquarium. Our results revealed significant differences between tadpole and adult microbiomes, as well as significant sex-dependent differences within the adult Wyoming toads, in terms of richness and composition. Thus, these findings have identified the baseline microbiome of this endangered species, and variables significantly influencing its composition. Ongoing studies of the only extant wild population are expected to identify taxa not present in captive toads, and potentially help design husbandry modifications to maximize survivability following reintroduction to the wild.

## Introduction

Amphibian species are critical to habitats due to their vital role spanning aquatic and terrestrial ecosystems, their role in the food web, their ability to serve as biological indicators, and their contributions to nutrient cycling, among others (Polasik et al., 2016). Thus, special efforts to protect amphibian species and ensure their survival are warranted.

At one time thought to be extinct in the wild, the Wyoming toad (*Anaxyrus baxteri*) is one of the most critically endangered North America amphibian species (Vincent & Abbott, 2015). The Wyoming toad populations were listed as endangered under the 1983 amendment to the Endangered Species Act (Lewis et al., 1985) and were thought to be extinct until a small population was discovered in Mortenson Lake in Albany County, Wyoming in 1987. In 1989, the last 10 remaining Wyoming toads were brought to the Cheyenne Mountain Zoo. The species was declared extinct in the wild in 1991, and the establishment of an *ex situ* breeding program followed in 1993 with the first reintroductions to their native range in 1995 (Lewis et al., 1985). By characterizing the captive microbiome of this species, we can potentially improve the microbial health of animals that are being produced for release so they may survive and subsequently breed *in situ*, increasing indigenous populations. Further indications to compare these findings with wild populations will improve our understanding of the impact of the microbiome on survivability after release. This information can facilitate reassessment of the way these animals are prepared for release to maximize survivability.

Previously, the Wyoming toad was found only in the Laramie Basin on floodplains of the Big and Little Laramie Rivers (southeast Wyoming) before irrigation practices changed the nature of the hydrology. The two highest contributing factors of population decline are *Batrachochytrium dendrobatidis* (Bd) infection and habitat destruction (Polasik et al., 2016). There is an association between the toad’s skin microbiome, serving as part of the host’s innate immune system, and resistance to Bd infection (Jiménez et al., 2019). This defensive microbiome relationship has also been seen in related species, including the boreal toad (*Anaxyrus boreas*) (Barnhart et al., 2017). Additionally, the environment in which individuals mature may have an important impact on the development of the animal’s microbiome in wild animal species across taxa (Comizzoli et al., 2021). Taken together, successful reintroductions of endangered animals may functionally involve both the species of interest, as well as that host-associated microbiomes of that species.

An animal’s microbiome is the collection of symbiotic microorganisms associated with that host organism. Microbiome dysbiosis can lead to negative host health impacts (immune, nutritional, and/or neurologic, for example) and compromised fitness (especially when facing changing environments). It is becoming clear that developmental establishment of a context-appropriate, beneficial microbiome is a large factor regulating the success of captive-bred species reintroduction conservation efforts (West et al., 2019).

There is concern among those working on these amphibian reintroduction projects that captive-bred individuals fail to develop the proper microbiota populations to withstand the context-specific stresses following release into native Wyoming habitats. In other words, these individuals may be ‘too sterile” or have fewer compatible microbiomes to survive and breed after reintroduction, possibly contributing to the inability of these populations to re-establish despite ongoing releases of these early life stage individuals. For example, in the year 2000 after 10,000 toads/tadpoles had been released, only 64 were seen in the wild (USGS, 2001). Little has been currently published about the healthy skin microbiome of the Wyoming toad, despite the theory that it could yield answers to some of the important conservation questions at hand. To that end, U. S. Fish and Wildlife Service (USFWS) and United States Geological Survey (USGS) have ongoing projects (2018-2023) assessing, in part, the *in situ* Wyoming toad microbiomes (A. Walters & Chalfoun, 2023). However, the microbiomes of toadlets produced at the breeding facilities are also unknown.

This project aimed to fill that knowledge gap and complement the ongoing Federal field microbiome research project. By detailing the microbiome of the captive Wyoming toad breeding population and the development of the offspring microbiome at Omaha’s Henry Doorly Zoo and Aquarium (OHDZA; Omaha, NE), our goal was to document the development of a baseline microbiome in those animals bred for release in Wyoming and compare to captive adult breeder population.

## Methods

### Toads

Samples were collected from immature (tadpole) and adult (approximately 1-4 years old) Wyoming toads (*Anaxyrus baxteri*) maintained at OHDZA. Toads are part of a captive colony housed in two isolation rooms (Room 2 and Room 8) located in OHDZA’s Amphibian Conservation Area. All adult breeder toads were group-housed in condos (*n* = 2 toads/condo) that allowed access to a constant filtered water bath as well as terrestrial substrate located in Room 2. Tadpoles were also housed in condos connected to the same filtered water system as in Room 2, which has never housed any other amphibian species except for the Wyoming toad. Additional tadpoles were housed in isolation Room 8, which was an empty isolation room prior to the successful spawning events of the 2022 breeding season. All water was from a filtered system in accordance with species regulations, to ensure 0ppm ammonia, <0.05ppm nitrite, <10ppm nitrate, and 68°F-75°F temperature.

Tadpoles were allowed to mature in Rooms 2 and 8 before being sent to various conservation sites in the Laramie Water Basin, WY for release later in the summer. A small portion of these tadpoles were kept back from release to be used as new breeding stock. Each room was kept in isolation and as a separate environment as to best fit the needs of the species housed there, with handlers changing PPE before entering each room and a work-flow order such that Wyoming toads were handled first by keeper staff throughout the day due to their conservation importance and susceptibility to disease as to not contaminate them with other species housed within the Amphibian Conservation Area. Protocol PPE includes lab coats, nitrile gloves, and rubber boots within the bio secure isolation room.

An Association of Zoos and Aquariums (AZA) Research Proposal form and protocols were submitted and approved by the OHDZA Animal Care and Use Committee in April 2022. As the animals were owned by the U.S. Fish and Wildlife Service, a Wyoming Toad Research Proposal was also submitted to the Wyoming Toad Recovery Team and approved by vote at their annual spring meeting (2022). Federal Permitting held under OHDZA allowed for housing and research on this species, in accordance with the Endangered Species Act.

### Sample collection

Mucosal samples were collected from adults using sterile cotton-tipped swabs. Each animal was maintained in a holding condo for the duration of sampling. Prior to any swabbing, they were rinsed front and back with approximately 100 mL of sterile water. Applying uniform pressure each time, the toad was swabbed while rotating the swab back and forth for 10 complete back-and-forth strokes in each of the following areas (in the order listed) for a total of 60 strokes per toad: along the right side of the abdomen, along the left side of the abdomen, along the ventral surface of the right hind leg, along the ventral surface of the left hind leg, along the spine (central dorsal surface) and along the mouth. Swabs were then broken off into sterile 1.5 mL Eppendorf tubes and stored on ice for transportation from the zoo to the lab (approximately 6 hours).

Cloacal swabs were collected from the adults following mucosal swabbing. A sterile cotton-tipped swab was advanced through the cloacal vent three times. The swab was then broken off into a sterile vial and stored on ice for transportation.

Due to the small surface area on the tadpoles, several tadpoles (five tadpoles in the first round of swabs, three in the second and third round of swabs) in the same cohort and living condo were swabbed with the same sterile cotton-tipped swab so as to yield a readable DNA sample while at the same time not stripping the animal of all their necessary mucosa. Animals were rinsed in a sterile water bath in group sizes as stated above, then moved to a transfer cup to have the water drained off as best as possible. Maintaining even pressure, the animals in each group were sampled with 10 complete back-and-forth strokes using a sterile swab. The swab tips were again broken off into sterile vials and the samples transported back to the laboratory on ice.

Biosecurity protocols put in place by the Wyoming Toad SSP and OHDZA’s Amphibian Conservation Area were followed for all Wyoming toad and tadpole handling, manipulating, and sampling. Each round of samples taken from within the Wyoming toad room required new PPE to be worn to ensure no/low pathogen transfer from external sources. Toads were handled one at a time, PIT tags verified for correct identities, and placed back into holding condos as soon as the swabs were complete. Tadpoles were removed from their main holding tanks to smaller volumes of water to facilitate sample collection. While in these smaller volumes of water, temperature remained constant and oxygen was provided from a remote, filtered oxygen source. Tadpole isolation ceased after one hour to ensure water quality parameters remained stable. Care was taken to minimize animal stress by collecting swabs as quickly as possible and closely monitoring the animals after swabbing to ensure their overall welfare remained of the highest quality. Any liquid or solid waste material was thoroughly disinfected with 70% ethanol and/or oxygenated powder bleach mixtures for 12 hours of contact time. Biological samples were stored appropriately and kept separate from any outside influences.

### DNA extraction

DNA was extracted using QIAamp PowerFecal Pro DNA extraction kits (Qiagen) according to the manufacturer instructions with the exception that DNA was eluted in 60 μL of EB buffer (Qiagen). DNA yields were quantified via fluorometry (Qubit 2.0, Invitrogen, Carlsbad, CA) using quant-iT BR dsDNA reagent kits (Invitrogen). Due to the low biomass, all DNA was used for 16S rRNA library preparation.

### 16S rRNA library preparation and sequencing

Library preparation and sequencing were performed at the University of Missouri (MU) Genomics Technology Core. Bacterial 16S rRNA amplicons were constructed via amplification of the V4 region of the 16S rRNA gene with universal primers (U515F/806R) previously developed against the V4 region, flanked by Illumina standard adapter sequences (Caporaso et al., 2011; W. A. Walters et al., 2011). PCR was performed as 50 μL reactions containing 100 ng metagenomic DNA, dual-indexed forward and reverse primers (0.2 μM each), dNTPs (200 μM each), and Phusion high-fidelity DNA polymerase (1U, Thermo Fisher). Amplification parameters were 98°C^(3 min)^ + [98°C^(15 sec)^ + 50°C^(30 sec)^ + 72°C^(30 sec)^] × 25 cycles + 72°C^(7 min)^. Amplicon pools were combined, mixed, and then purified by addition of Axygen Axyprep MagPCR clean-up beads to an equal volume of 50 μL of amplicons and incubated for 15 minutes at room temperature. Products were washed multiple times with 80% ethanol and the pellet was resuspended in 32.5 μL EB buffer (Qiagen), incubated for two minutes at room temperature, and then placed on the magnetic stand for five minutes. The final amplicon pool was evaluated using an Advanced Analytical Fragment Analyzer automated electrophoresis system, quantified using quant-iT HS dsDNA reagent kits, and diluted according to the Illumina standard protocol for sequencing as 2×250 bp paired-end reads on the MiSeq instrument.

### Bioinformatics

16S rRNA sequences were processed using Quantitative Insights Into Microbial Ecology 2 (QIIME2) v2021.8 (Bolyen et al., 2019). Illumina adapters and primers were trimmed from forward and reverse reads with cutadapt (Martin, 2011). Reads were then truncated to 150 base pairs, then denoised into unique amplicon sequence variants (ASVs) using DADA2 (Callahan et al., 2016). Unique sequences were then assigned taxonomy using an a sklearn algorithm and the QIIME2-provided 99% non-redundant SILVA v138 reference database (Quast et al., 2013) trimmed to the 515F/806R (Caporaso et al., 2011) region of the 16S rRNA gene. Alpha diversity metrics (Chao-1 and Simpson Indices) were determined using the *microbiome* (Lahti & Shetty, 2017) and *vegan* (Oksanen et al., 2014) libraries. Differences in beta diversity were visualized with principal coordinate analysis (PCoA) using Bray-Curtis (weighted) distances. Briefly, a distance matrix was generated with the *vegdist* function from the *vegan* library using a quarter-root transformed feature table. PCoAs were performed using the *ape* (Paradis & Schliep, 2019) library with a Calliez correction. The cladogram was generated using Graphlan v1.1.4 (Asnicar et al., 2015).

### Statistics

Univariate data (reported as mean ± SE) were first tested for normality using the Shapiro-Wilk method, followed by the appropriate parametric or non-parametric test. Whenever possible and practical, multifactor tests (e.g., two-way analysis of variance, ANOVA) were used. Differences in multivariate data were tested using permutational multivariate ANOVA (PERMANOVA) and visualized using PCoAs. PERMANOVAs were performed with 9,999 permutations. Both PERMANOVA and PCoA were performed using weighted (Bray-Curtis) distances. Differential abundance testing was performed using analysis of composition of microbiomes with bias correction 2 (ANCOM-BC2) with a significance threshold of a Benjamani-Hochberg-corrected *p* < 0.05 (Benjamini & Hochberg, 1995; Lin & Peddada, 2020). Structural zeroes (i.e., taxa present in ≥ 1 group and absent in in ≥ 1 group) were also reported. Pairwise ANCOM-BC2 was utilized as necessary. Microbial community analysis was performed using the open-source R statistical software v4.2.2 (Team, 2010).

## Results

### Sequencing data pass quality control

All but two samples resulted in usable data, with a total of 8,945,498 high-quality sequence reads, and a mean of 95,165 ± 7,297 sequence reads per sample. Samples from adults returned greater sequence read counts than samples from tadpoles (mean ± SE = 129,198 ± 10,886 in adults, 59,652 ± 6,391 in tadpoles, *p* < 0.001, Mann-Whitney rank sum test). Testing for group differences in sequencing depth separately within adults and tadpoles failed to detect any significant differences, although there was a trend toward greater sequence read counts in samples from male adults compared to female adults (**Supplemental Figure 1A**, *p* = 0.086, F = 3.0, two-way ANOVA) as well as a trend towards greater reads in mucosal samples compared to cloacal samples (*p* = 0.095, F = 2.9, two-way ANOVA). No difference in read counts were observed between tadpole colonies (**Supplemental Figure 1B**).

### Amplicon sequence variant (ASV) coverage

Denoising paired-end sequences into amplicon sequence variants (ASVs) recovered an average of 62,095 ± 5,824 features per sample ranging from 140 to 229,727 features per sample. As expected from the aforementioned increase in sequencing coverage, more ASVs were detected in adult samples compared to tadpole samples (mean ± SE; 90,908 ± 7,827 in adults, 32,030 ± 6,093 in tadpoles, *p* < 0.001, Mann-Whitney rank sum test). No significant differences in ASV counts were detected between groups within each age. Given the nature of low biomass samples (i.e., swabs) yielding low feature counts, we estimated the achieved sampling of each community using Good’s coverage. Each sample yielded a Good’s coverage of greater than 99.9%, thus the full, unrarefied feature table (ASV counts per sample) was used in all further analyses.

### Adult and tadpole Wyoming toad microbiomes differ in composition

An initial survey of the entire dataset revealed significant age-associated differences in richness (*p* < 0.001, Mann-Whitney rank sum test), although these were difficult to interpret in the context of the aforementioned difference in sequencing depth. Within adults, samples collected from mucosal tissues exhibited increased community richness relative to cloacal samples (**Figure 1A**, *p* = 0.011, F = 7.0, two-way ANOVA). No sex-dependent effects on sample richness were observed (*p* = 0.325, F = 0.99, two-way ANOVA). Samples collected from adult males exhibited increased alpha diversity (Simpson Index) relative to females with no effect of sample type (**Figure 1B**, Sex: *p* = 0.016, F = 6.3; Sample Type: *p* = 0.59, F = 0.29, two-way ANOVA). Within tadpole samples, significant differences in community richness were observed between colonies (Room:Colony: *p* = 0.005, F = 6.1, nested two-factor ANOVA). Within Room 2, samples collected from Colony F exhibited increased richness compared to Colony A (**Figure 1C**, *p* = 0.033, Tukey *post hoc*). No significant differences in alpha diversity were detected between rooms or colonies (**Figure 1D**).

**Figure 1.**
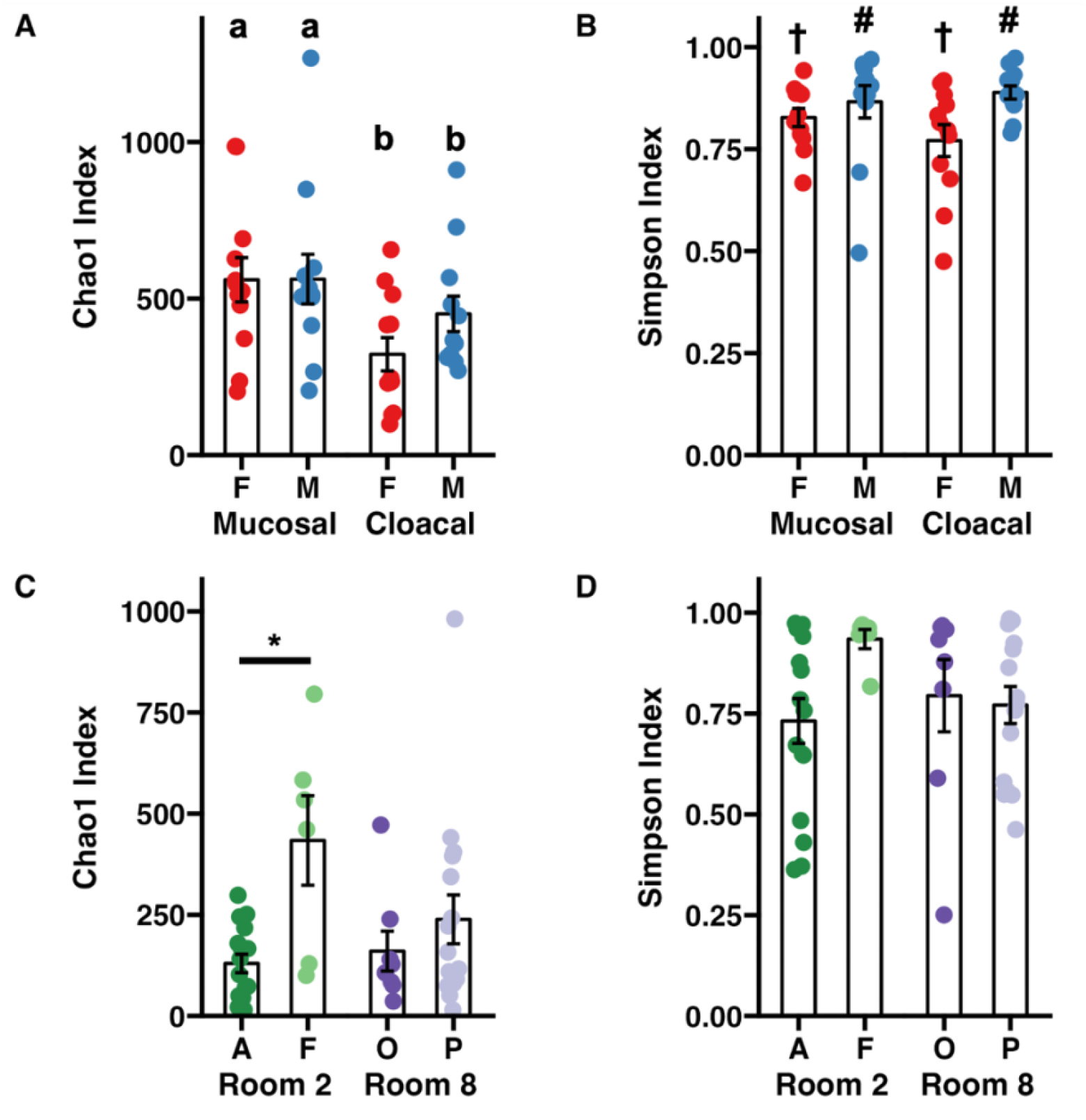
Dot plots depicting adult Wyoming toad (**A**) richness and (**B**) diversity and tadpole (**C**) richness and (**D**) diversity. Differences between adult samples were assessed using a two-way ANOVA. Differing letters represent sex-dependent differences of p < 0.05. Differing symbols represent sample type-dependent differences of p < 0.05. Differences between tadpole samples were assessed using a nested two-factor ANOVA. Pairwise differences between tadpole colonies were assessed using a Tukey post hoc test. * p < 0.05.

Visualization of beta diversity among the entire dataset using principal coordinate analysis (PCoA) with weighted (Bray-Curtis) distances showed complete separation of samples from adults and tadpoles along the first principal coordinate (**Figure 2**). One-way PERMANOVA comparing adult and tadpole samples using Bray-Curtis distances confirmed a significant difference in community composition between age groups (*p* < 0.001, F = 22.98). We then identified differentially abundant phyla between adults and tadpoles using analysis of composition of microbiomes with bias correction (ANCOM-BC2). Of the 35 detected phyla, 12 were differentially abundant including *Actinobacteriota* and *Patescibacteria* enriched in adults and WPS-2 and *Bdellovibrionota* enriched in tadpoles. The phyla *Synergistota* and *Fermentibacterota* were only detected in adults whereas *Hydrogenedentes, Latescibacterota, Modulibacteria*, and *Halanaerobiaeota* were only detected in tadpoles (**Supplemental File 1**). Collectively, these data demonstrate large age-dependent differences in alpha and beta diversity as well as unique taxonomic signatures within each group. We next stratified the data by age group to better assess the influence of sex and sample site in the adult toads, and room and cohort in the tadpoles.

**Figure 2.**
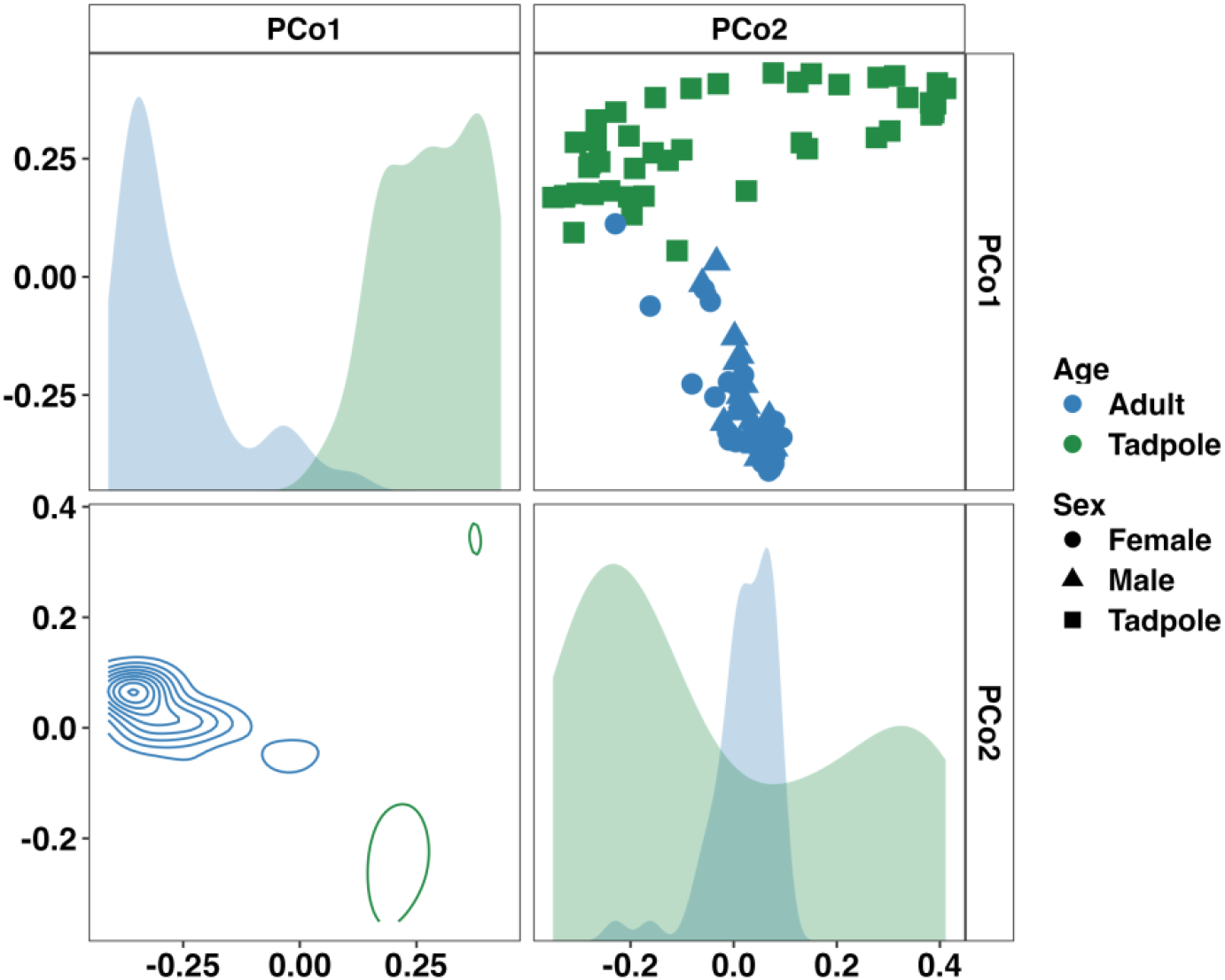
Principal coordinate analysis matrix using weighted Bray-Curtis distances depicting significant differences in community composition between adult and tadpole Wyoming toads along the first (22.75%) and second (9.06%) principal coordinates. F = 23.0, p < 0.001. One-way PERMANOVA.

### Adult Wyoming toads harbor sex-specific microbiomes, on mucosa and cloaca

Considering first the samples from adults, a sex- and sample type-dependent effect on beta-diversity was observed (Sex: *p* < 0.001, F = 3.55; Sample Type: *p* = 0.014, F = 1.92, two-way PERMANOVA). Visualizing these communities using a PCoA revealed a sex-dependent separation along the third principal coordinate (**Figure 3**). Differential abundance testing using ANCOM-BC2 found only *Armatimonadota* to be significantly enriched in males (**Supplemental Figure 2**). Additionally, *Fermentibacterota* and *Fibrobacterota* were only detected in males whereas *Nitrospirota* was only observed in females. Only one, unresolved bacterial taxa was differentially abundant between mucosal and cloacal samples. At the family level, 41 taxa were differentially abundant between males and females (22 and 19 taxa, respectively). Many families within the phylum *Pseudomonadota* including *Alcaligenaceae, Labraceae*, and *Aeromonadaceae* were enriched in males. *Deinococcaceae* (Phylum Deinococcota), *Nitrospiraceae* (Phylum *Nitrospirota*), and multiple families within the phylum Cyanobacteria were enriched in females (**Supplemental Figure 2**). Only three families were differentially abundant between mucosal and cloacal samples. *Mycobacteriaceae* and *Solirubrobacteraceae* (phylum *Actinobacteriota*) and one unresolved bacteria were enriched in mucosal samples. An additional 114 families were found only in males (53) or females (61). A full list of differentially abundant taxa at the phylum and family levels in adults is provided in **Supplemental File 2**. These data demonstrate large sex-dependent differences in the adult Wyoming toad microbiome.

**Figure 3.**
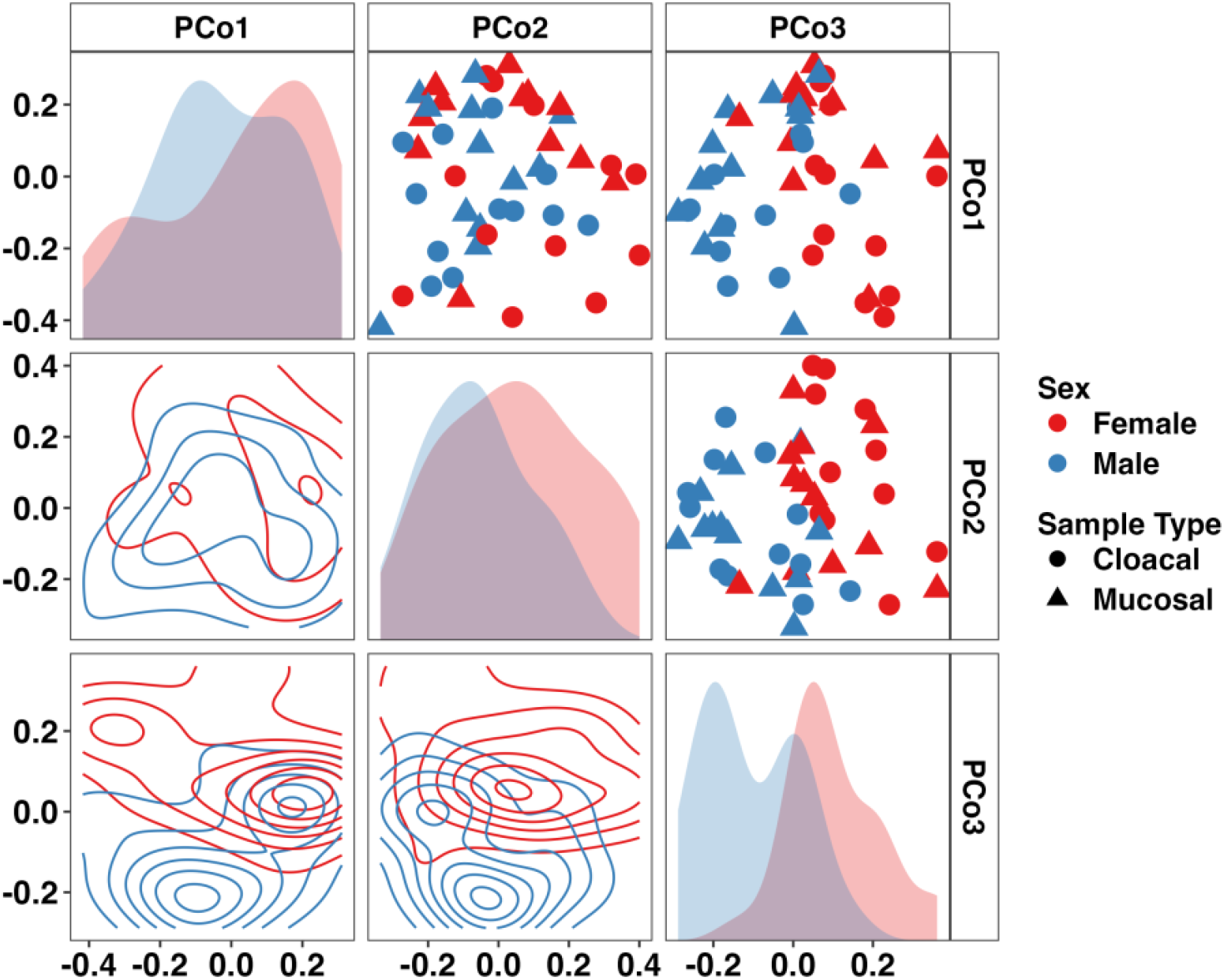
Principal coordinate analysis matrix using weighted Bray-Curtis distances depicting differences in community composition between sex and sample type in adult Wyoming toads along the first (16.31%), second (13.45%), and third (9.85%) principal coordinates. Sex- (p < 0.001, F = 3.6) and sample type-dependent (p = 0.014, F = 1.9) effects on community composition were observed. Two-way PERMANOVA.

### Room-specific husbandry subtly affects the Wyoming toad tadpole microbiome

Focusing on the separate tadpole colonies, we first identified significant differences in the weighted microbial composition between colonies but not rooms overall (**Figure 4**, Room: *p* = 0.053, F = 1.68; Room:Colony: *p* = 0.021, F = 1.62, nested two-factor PERMANOVA). Pairwise comparisons revealed significant, albeit modest, differences in community composition between Colony A-Colony F (*p* = 0.044, F = 1.83) and Colony F-Colony P (*p* = 0.010, F = 2.13). We again applied ANCOM-BC2 at the phylum and family levels to identify differentially abundant taxa within all four colonies. No significant differentially abundant taxa were identified at the phylum or family levels. Twenty of the thirty-three resolved phyla were observed in all four tadpole colonies. Using pairwise ANCOM-BC2 comparisons, five families were identified as enriched predominantly in Colony F. These families included an uncultured *Rhodospirillales* and uncultured *Alphaproteobacteria* (phylum *Pseudomonadota*), uncultured RBG-13-54-9 (phylum *Chloroflexi*), *Sporichthyaceae* (phylum *Actinobacteriota*), and *Pedosphaeraceae* (phylum *Verrucomicrobiota*). A full list of differentially abundant taxa at the phylum and family levels in each tadpole colony is provided in **Supplemental File 3**.

**Figure 4.**
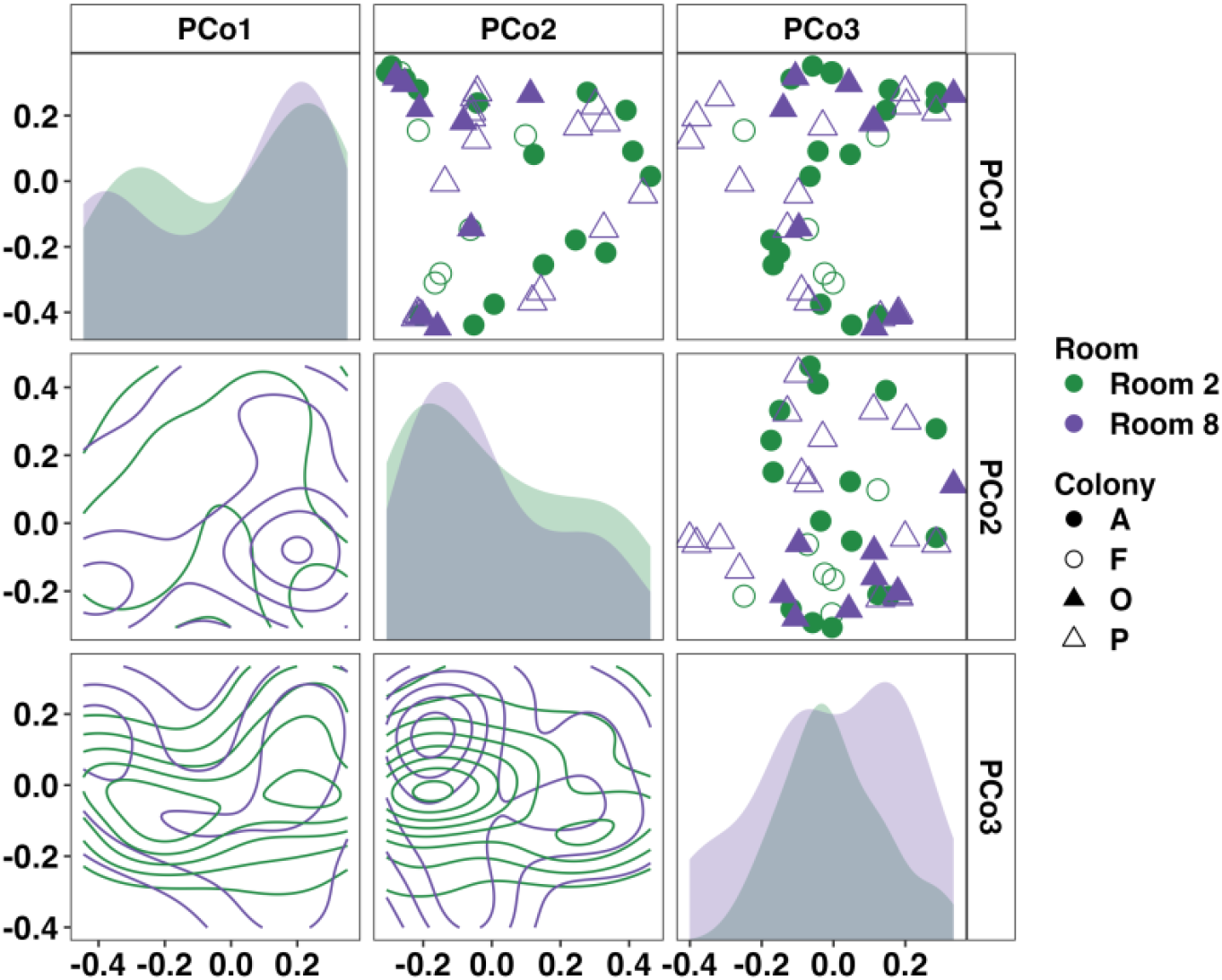
Principal coordinate analysis matrix using weighted Bray-Curtis distances depicting differences in community composition between Wyoming tadpole colonies along the first (19.41%), second (13.43%), and third (7.89%) principal coordinates. Significant colony-dependent differences were observed. Room: *p* = 0.053, F = 1.7; Room:Colony: *p* = 0.021, F = 1.6. Nested two-factor PERMANOVA.

### Longitudinal analysis of adult and tadpole Wyoming toads

To characterize the stability of the Wyoming toad microbiome, samples were collected at three separate time points for all adults and 2-3 separate time points for tadpole colonies. Time points 1 and 2 were collected 15 days apart and time points 2 and 3 were collected 24 days apart. Given the modest difference in beta diversity between sample types in adults, we combined cloacal and mucosal samples for longitudinal analysis in adults. Two-way PERMANOVA analysis of adult samples revealed no significant effect of time on microbial composition (Sex: *p* < 0.001, F = 3.6; Time Point: *p* = 0.123, F = 1.3). Pairwise comparison of every adult sample across time using Bray-Curtis distances supported this as no patterns of dissimilarity were revealed (**Figure 5A**). When assessing tadpoles, two-way PERMANOVA analysis revealed significant differences in community composition between colonies (*p* < 0.001, F = 2.1), across time (*p* < 0.001, F = 5.0), and an interaction of the two (*p* = 0.001, F = 1.7), however, visualization of the pairwise Bray-Curtis distances revealed high dissimilarity between all samples with no discernable patterns (**Figure 5B**). The dramatic morphological changes occurring between sampling periods in the Wyoming toad tadpole may contribute to the longitudinal shifts in microbiome composition (**Supplementary Figure 3)**.

**Figure 5.**
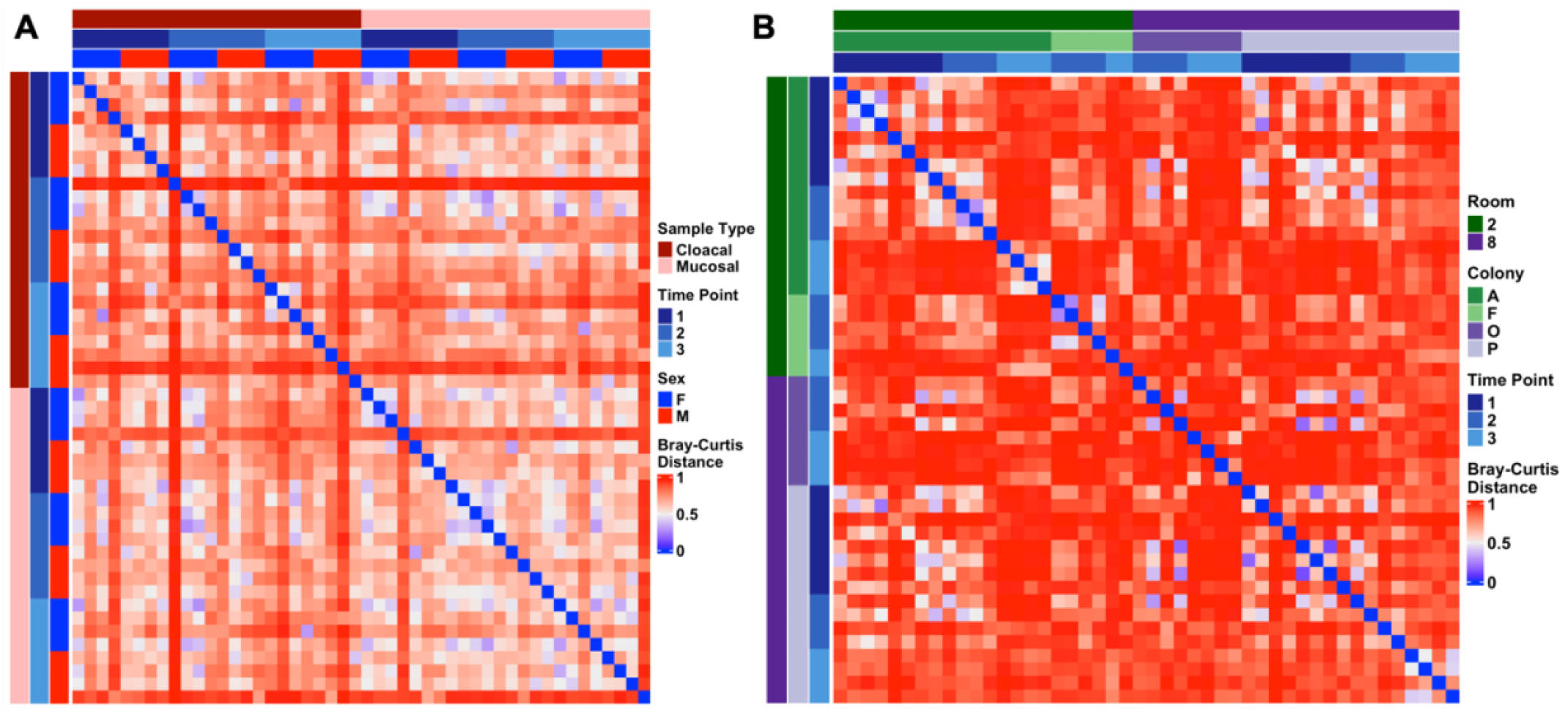
Heatmaps depicting longitudinal Bray-Curtis distances within (**A**) adult and (**B**) tadpole samples. Metadata describing sample type and timepoint are displayed as colored bars along the left and top of each heatmap.

## Discussion

As expected, the microbiomes of adult breeding animals differed in richness and alpha-diversity from that of the tadpoles, suggesting the dynamic establishment of the mucosal microbiome in amphibian species. This is of importance to the *ex situ* breeding programs, as many of these tadpoles are being released back into the wild ranges at such young ages that the mucosal microbiome may not have been established prior to release. While this may have the benefit of allowing the animal to develop a microbiome in accordance with their environmental stressors encountered post-release, there is also the potential that these animals are not necessarily strong enough to support survivability when faced with natural stressors. Further indications may include a more detailed study to determine the timepoint in the immature animal’s life to fully appreciate the effects of this phenomenon.

Sex-specific microbiome of the adult breeding toads, both in cloaca and mucosal sampling, appears to be novel within the literature of amphibian species. The animals in this study were sampled during the breeding season and females had been on an artificial hormone therapy, so it is unclear at this time the role this might play in our results. While not directly related, a study on the cloacal microbiome of free-living rufous-collared sparrows *(Zonotrichia capensis)* differed greatly among the sexes during the breeding season (Escallón et al., 2019), showing a similar phenomenon in avian species. Further study may be indicated to tease out the biological relevance of sex-dependent diversity in amphibians.

Our results from captive animals could be further leveraged by comparing the microbiomes of those surviving animals at the Laramie Basin release sites to decide if husbandry modifications are necessary to propagate tadpoles and toadlets with proper survivable microbiome populations. To our knowledge, there has not been a study done in related species that would compare side by side toad species raised in captivity with their wild ranging counterparts. Such a study would answer the question raised by caretakers that these animals appear too sterile for hard releases. Additionally, there is a potential for inoculation studies or supplementation of the deficits in the captive raised tadpoles should the need seem fit. This would allow maximization of the survivability of the toadlets once they are released back into their natural environments by maximizing the potential for their defensive microbiome, providing them with the microbial populations that are necessary to inhabit the wetlands of their native ranges.

Going beyond the Wyoming toad populations, it is important to note that *Batrachochytrium dendrobatidis* (*Bd*) infection is the leading cause of rapid decline of amphibian species worldwide (Jiménez et al., 2019). If there is an advantage to the skin microbiome, this information might not only help with the *ex situ* breeding program in this species, but also benefit other species facing similar challenges. Further analysis into how the microbial species discovered in this project interact with *Bd* could yield further insight into answering the question of susceptibility to such infection. Since a defensive microbiome has been seen in related species, *Anaxyrus boreas* (Barnhart et al., 2017), further detailing of these advantages may reveal insights into mitigating such a devastating disease.

## Acknowledgements

We would like to thank the staff at Omaha’s Henry Doorly Zoo and Aquarium’s Amphibian Conservation center for their support of this project, including J. Krebs. We would also like to thank the keeper staff at the Amphibian Conservation Center for their role in sample collection and ensuring animal husbandry and welfare needs were met. We thank the University of Missouri College of Veterinary Medicine’s Veterinary Research Scholars Program (VRSP) for program support and mentorship. Stipend support for Nicole Scarberry was provided by the Morris Animal Foundation through the Veterinary Student Scholars grant.

## Data availability statement

All sequencing data and associated metadata have been deposited in the NCBI Sequence Read Archive and are freely available BioProject ID <u>PRJNA943179</u>.

### Author Contributions

JH, BM, and ACE conceptualized the project. JH, BM, ACE, and NS designed the study. NS, DH, and JH collected the data. NS, ZM, and ACE interpreted the data. NS and ZM drafted the manuscript. All authors contributed feedback on manuscript revisions.

## Figures and figure legends

**Supplementary Figure 1.**
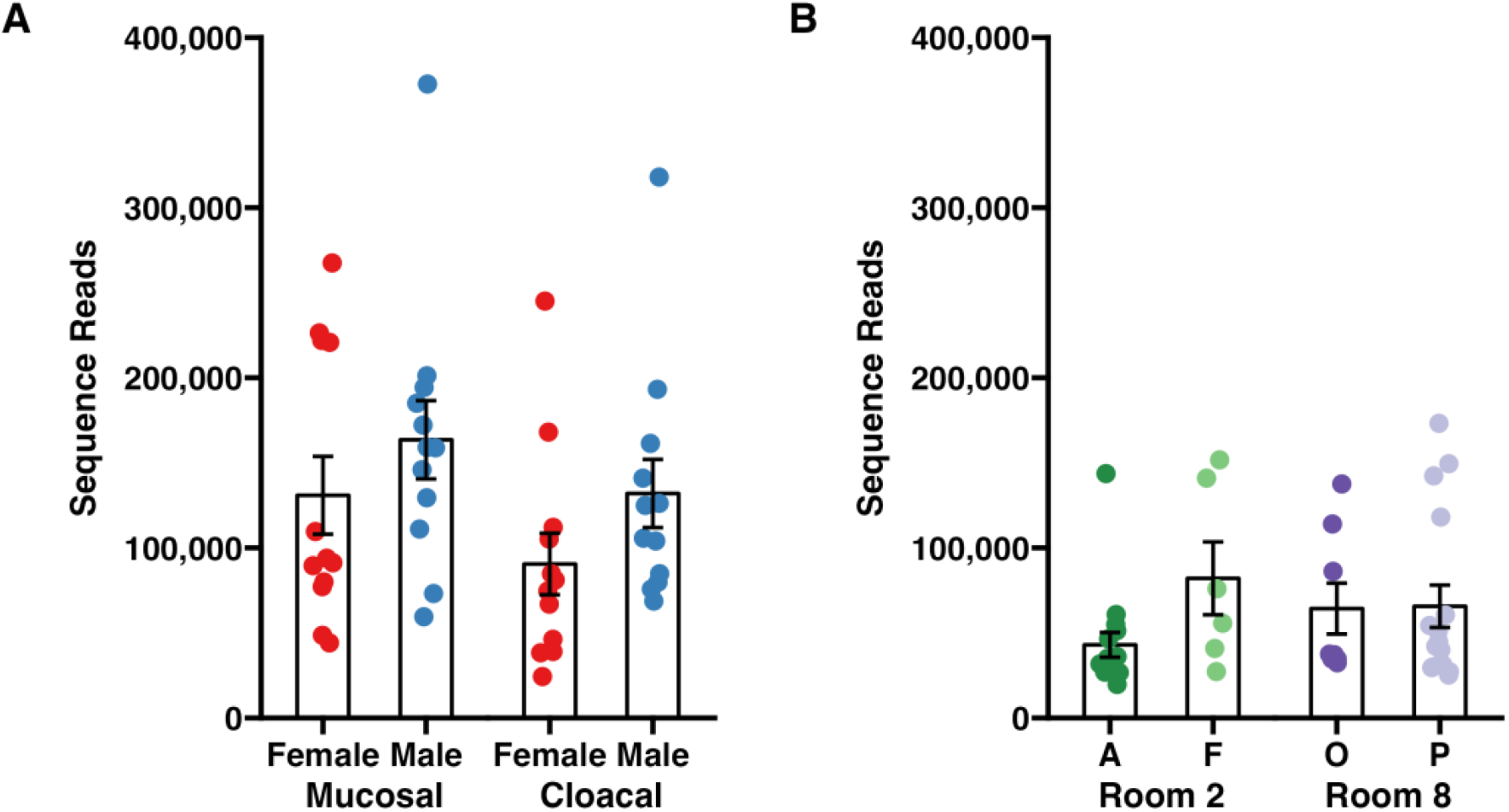
Forward read counts recovered from 16S rRNA sequencing of **(A)** adult and **(B)** tadpole Wyoming toad samples. Differences between adult samples were assessed using a two-way ANOVA. Sex: *p* = 0.086, F = 3.1; Sample Type: *p* = 0.095, F = 2.9. Differences between tadpole samples were assessed using a nested two-factor ANOVA. Room: *p* = 0.364, F = 0.84; Room:Colony: *p* = 0.172, F = 1.8.

**Supplementary Figure 2.**
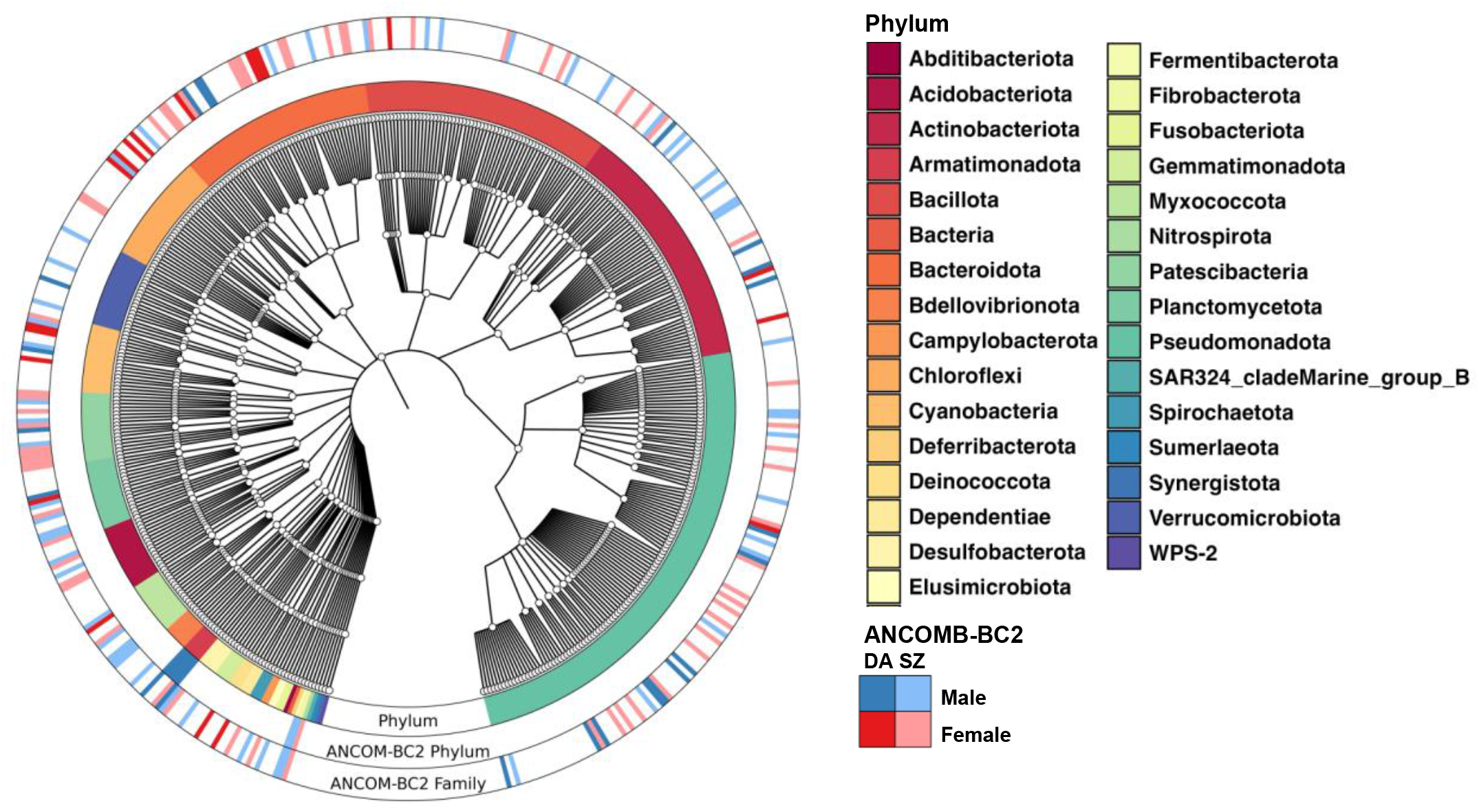
Family-level cladogram of taxa resolved in adult samples. Rings represent (from innermost out) Phylum-level classification, differentially abundant phyla, and differently abundant families between sexes. DA: Differentially abundant by significance (Benjamini-Hochberg corrected *p* < 0.05). SZ: Structural zero determined by presence/absence. ANCOM-BC2.

**Supplementary Figure 3.**
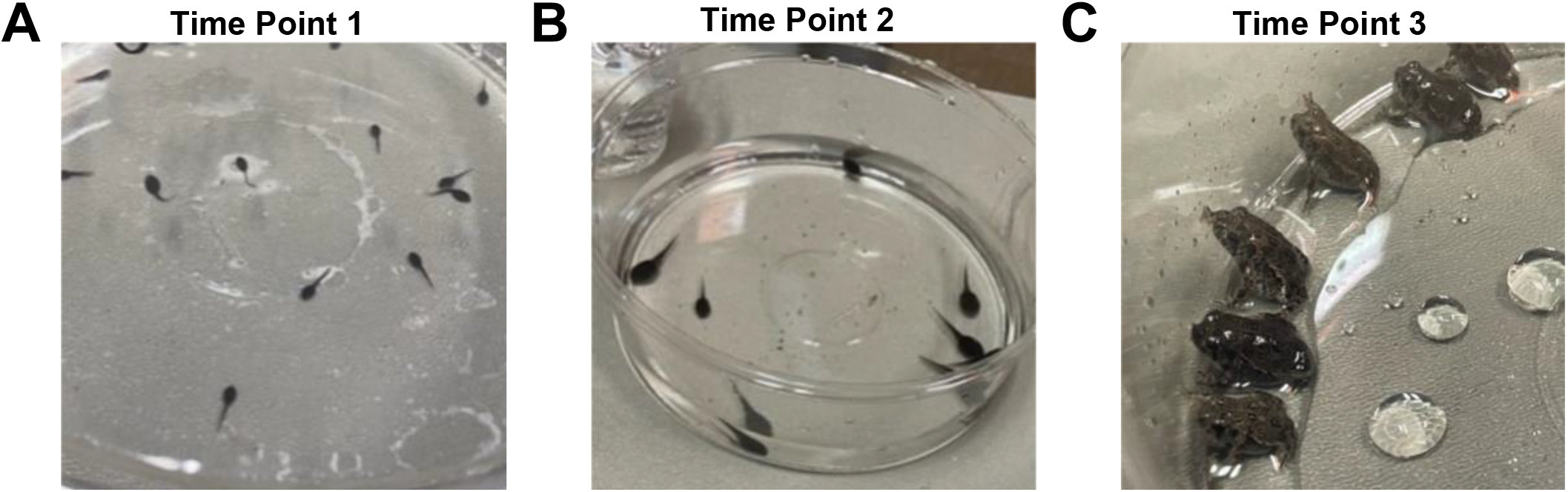
Representative images of Wyoming toad tadpoles at the three collection timepoints. **(A) Time Point 1:** 1-5 days post hatch. **(B) Time Point 2:** Midpoint Evaluation **(C) Time Point 3:** 2 days post metamorphosis.

## References

Asnicar, F., Weingart, G., Tickle, T. L., Huttenhower, C., & Segata, N. (2015). Compact graphical representation of phylogenetic data and metadata with GraPhlAn. PeerJ, 3, e1029. https://doi.org/10.7717/peerj.1029

Barnhart, K., Forman, M. E., Umile, T. P., Kueneman, J., McKenzie, V., Salinas, I., Minbiole, K. P. C., & Woodhams, D. C. (2017). Identification of Bufadienolides from the Boreal Toad, Anaxyrus boreas, Active Against a Fungal Pathogen. Microbial Ecology, 74(4), 990–1000. https://doi.org/10.1007/s00248-017-0997-8

Benjamini, Y., & Hochberg, Y. (1995). Controlling the False Discovery Rate: A Practical and Powerful Approach to Multiple Testing. Journal of the Royal Statistical Society: Series B (Methodological), 57(1), 289–300. https://doi.org/10.1111/j.2517-6161.1995.tb02031.x

Bolyen, E., Rideout, J. R., Dillon, M. R., Bokulich, N. A., Abnet, C. C., Al-Ghalith, G. A., Alexander, H., Alm, E. J., Arumugam, M., Asnicar, F., Bai, Y., Bisanz, J. E., Bittinger, K., Brejnrod, A., Brislawn, C. J., Brown, C. T., Callahan, B. J., Caraballo-Rodriguez, A. M., Chase, J., … Caporaso, J. G. (2019). Reproducible, interactive, scalable and extensible microbiome data science using QIIME 2. Nat Biotechnol, 37(8), 852–857. https://doi.org/10.1038/s41587-019-0209-9

Callahan, B. J., McMurdie, P. J., Rosen, M. J., Han, A. W., Johnson, A. J., & Holmes, S. P. (2016). DADA2: High-resolution sample inference from Illumina amplicon data. Nat Methods, 13(7), 581–583. https://doi.org/10.1038/nmeth.3869

Caporaso, J. G., Lauber, C. L., Walters, W. A., Berg-Lyons, D., Lozupone, C. A., Turnbaugh, P. J., Fierer, N., & Knight, R. (2011). Global patterns of 16S rRNA diversity at a depth of millions of sequences per sample. Proc Natl Acad Sci U S A, 108 Suppl 1(upplement_1), 4516–4522. https://doi.org/10.1073/pnas.1000080107

Comizzoli, P., Power, M. L., Bornbusch, S. L., & Muletz-Wolz, C. R. (2021). Interactions between reproductive biology and microbiomes in wild animal species. Animal Microbiome, 3(1), 87. https://doi.org/10.1186/s42523-021-00156-7

Escallón, C., Belden, L. K., & Moore, I. T. (2019). The Cloacal Microbiome Changes with the Breeding Season in a Wild Bird. Integrative Organismal Biology, 1(1), oby009. https://doi.org/10.1093/iob/oby009

Jiménez, R. R., Alvarado, G., Estrella, J., & Sommer, S. (2019). Moving Beyond the Host: Unraveling the Skin Microbiome of Endangered Costa Rican Amphibians. Frontiers in Microbiology, 10, 2060. https://doi.org/10.3389/fmicb.2019.02060

Lahti, L., & Shetty, S. (2017). Tools for microbiome analysis in R. Version 2.1.26. http://microbiome.github.com/microbiome

Lewis, D. L., Baxter, G. T., Johnson, K. M., & Stone, M. D. (1985). Possible Extinction of the Wyoming Toad, Bufo hemiophrys baxteri. Journal of Herpetology, 19(1), 166–168. https://doi.org/10.2307/1564434

Lin, H., & Peddada, S. D. (2020). Analysis of compositions of microbiomes with bias correction. Nat Commun, 11(1), 3514. https://doi.org/10.1038/s41467-020-17041-7

Martin, M. (2011). Cutadapt removes adapter sequences from high-throughput sequencing reads. Embnet Journal, 17.

Oksanen, J., Blanchet, F. G., Kindt, R., Legendre, P., Minchin, P. R., O’Hara, R. B., Simpson, G. L., Solymos, P., Stevens, M. H. H., & Wagner, H. (2014). Vegan: Community Ecoloy Package. R package version 2.2-0.

Paradis, E., & Schliep, K. (2019). ape 5.0: an environment for modern phylogenetics and evolutionary analyses in R. Bioinformatics, 35(3), 526–528. https://doi.org/10.1093/bioinformatics/bty633

Polasik, J. S., Murphy, M. A., Abbott, T., & Vincent, K. (2016). Factors limiting early life stage survival and growth during endangered Wyoming toad reintroductions. The Journal of Wildlife Management, 80(3), 540–552. https://doi.org/10.1002/jwmg.1031

Quast, C., Pruesse, E., Yilmaz, P., Gerken, J., Schweer, T., Yarza, P., Peplies, J., & Glockner, F. O. (2013). The SILVA ribosomal RNA gene database project: improved data processing and web-based tools. Nucleic Acids Res, 41D1), D590–6. https://doi.org/10.1093/nar/gks1219

Team, R. D. C. (2010). R: A Language and Environment for Statistical Computing. http://www.R-project.org

USGS. (2001). Population and habitat viability assessment for the Wyoming toad (Bufo baxteri): Final workshop report (p. 109) [Report]. Conservation Breeding Specialist Group; USGS Publications Warehouse. http://pubs.er.usgs.gov/publication/70159722

Vincent, K., & Abbott, T. (2015). First Revised Recovery Plan for Wyoming Toad.

Walters, A., & Chalfoun, A. (2023). Research and Monitoring of Wyoming Toad Reintroductions: Linking Survival, Behavior and Genetics to Inform Species Recovery. https://www1.usgs.gov/coopunits/project/170219780096/awalter8

Walters, W. A., Caporaso, J. G., Lauber, C. L., Berg-Lyons, D., Fierer, N., & Knight, R. (2011). PrimerProspector: de novo design and taxonomic analysis of barcoded polymerase chain reaction primers. Bioinformatics, 27(8), 1159–1161. https://doi.org/10.1093/bioinformatics/btr087

West, A. G., Waite, D. W., Deines, P., Bourne, D. G., Digby, A., McKenzie, V. J., & Taylor, M. W. (2019). The microbiome in threatened species conservation. Biological Conservation, 229, 85–98. https://doi.org/10.1016/j.biocon.2018.11.016

